# Full life-cycle models from ring-recovery data: estimating fecundity from age ratios at capture

**DOI:** 10.1101/352658

**Authors:** Todd. W. Arnold

**Author notes:** Twitter: @Todd_W_Arnold.

## Abstract

1. Tag-recovery data from organisms captured and marked post breeding are commonly used to estimate juvenile and adult survival. If annual fecundity could also be estimated, tagging studies such as European and North American bird-ringing schemes could provide all parameters needed for building full life-cycle projection models.
2. I modified existing tag-recovery models to allow estimation of annual fecundity using age composition and recapture probabilities obtained during routine banding operations of northern pintails (*Anas acuta*) and dark-eyed juncos (*Junco hyemalis*), and I conducted simulations to assess estimator performance in relation to sample size.
3. For pintails, population growth rate from band-recovery data (λ = 0.929, SD 0.060) was similar but less precise than count-based estimates from the Waterfowl Breeding Pair and Habitat Survey (λ 0.945, SE 0.001). Models with temporal variation in vital rates indicated that annual population growth in pintails was driven primarily by variation in fecundity. Juncos had lower survival but greater fecundity, and their estimated population growth rate (λ 1.01, SD 0.19) was consistent with count-based surveys (λ 0.986).
4. Simulations indicated that reliable (CV < 0.10) estimates of fecundity could be obtained with >1000 same-season live encounters. Although precision of survival estimates depended primarily on numbers of adult recoveries, estimates of population growth rate were most sensitive to total number of live encounters.
5. *Synthesis and applications*: Large-scale ring-recovery programmes could be used to estimate annual fecundity in many species of birds, but the approach requires better data curation, including accurate assessment of age, better reporting of banding totals and greater emphasis on obtaining and reporting same-season live encounters.

## 1 | INTRODUCTION

Tag-recovery (a.k.a. ring- or band-recovery) models are widely used to estimate annual survival using data on numbers of individuals surviving different intervals between tagging and reported time of death (Seber 1970; Brownie *et al.* 1978). Unlike live encounter data from restricted study areas, which provide estimates of apparent survival φ = (1 – mortality)*(1 – permanent emigration), dead recovery data can provide estimates of true survival *S* = (1 – mortality) provided that tag recoveries occur from throughout the population’s potential dispersal or migratory range. This occurs most commonly with harvested populations of birds and fish (Brownie *et al.* 1978; Pollock, Hearn & Polacheck 2002), although dead-recovery models have also been applied to unharvested species (Francis 1995; Siriwardena, Baillie & Wilson1998). Dead recoveries can also be combined with live-encounter data from restricted study areas to estimate true survival and permanent emigration (Burnham 1993; Barker 1997).

If individuals can be reliably assigned to age classes at the time of marking, tag-recovery models can be used to estimate age-specific survival (Seber 1971; Brownie *et al.* 1978; Pollock, Hearn & Polacheck 2002). Most typically with birds, this approach has been used to provide age-specific survival estimates for juveniles (*S*_*j*_) and adults (*S*_*a*_), but it can also be used for three or more age classes provided age classes can be recognized at marking (Brownie *et al.* 1978). For monogamous species that reach sexual maturity as one-year-olds and have limited sex- or age-specific variation in survival or fecundity (e.g., many small birds and mammals), population dynamics can be modeled using a simple one-stage projection model that captures most of the important variation in vital rates:

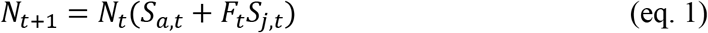
where *S*_*a,t*_ + *F*_*t*_*S*_*j,t*_ is the population growth rate, λ_*t*_ = *N*_*t*+1_/*N*_*t*_. Thus, tag-recovery models provide everything needed to estimate λ_*t*_ except annual fecundity *F*_*t*_.

Fecundity can be estimated using age ratios (*N*_*j,t*_/*N*_*a,t*_) collected during post birth-pulse surveys, and age ratios are commonly used when juveniles and adults can be readily distinguished during survey counts (Harris, Kauffman & Mills 2007; Weegman *et al.* 2016). Wildlife managers have long used age ratios of harvested individuals (*H*_*j,t*_/*H*_*a,t*_) to measure annual fecundity, but because juveniles are often more vulnerable to harvest than adults, tag recovery data are needed to adjust these data for relative vulnerability to harvest:

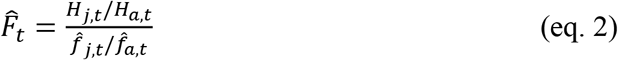
where 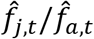 is the ratio of juvenile to adult harvest rates (Zimmerman *et al.* 2010). Age ratios at capture can provide similar estimates of population-level fecundity (Specht & Arnold 2018), but if capture methods are biased towards juveniles, fecundity estimates will be positively biased. However, live recaptures during the initial banding period could be used to assess age-specific vulnerability to capture and estimate the true underlying age distribution, similarly to vulnerability-adjusted age ratios at harvest (Zimmerman *et al.* 2010). Even if estimation of capture vulnerability is not possible, age ratios at capture might nevertheless provide a reliable index of annual fecundity. Although uncorrected age ratios at capture have been used to assess population-level fecundity (Mazerolle *et al.* 2005; Ross *et al.* 2017; Specht & Arnold 2018), models to estimate fecundity from initial capture data have not been formally developed for tag-recovery studies, although there are close equivalents in the live-encounter literature (Pradel 1996; Link & Barker 2005).

My objectives in this paper are to develop and apply population projection models including fecundity, juvenile survival and adult survival derived solely from tagging data (i.e., age-specific counts of numbers of birds banded during post-season capture occasions, recaptured alive during the same season as originally marked or subsequently found dead any time after marking). I apply these models to two species of North American birds. Northern pintails (*Anas acuta*) have experienced a prolonged population decline and previous studies have shown it cannot be explained by declining survival (Bartzen & Dufour 2017), suggesting that lowered fecundity may be the cause (Specht & Arnold 2018), but to date there have been no integrated analyses for pintails to identify relative contributions of different vital rates to observed population changes (Koons, Arnold & Schaub 2017). Dark-eyed juncos (*Junco hyemalis*) are a widespread passerine that has been well-studied at several localized and primarily southern breeding sites (Nolan *et al.* 2002), but most of their breeding range occurs in remote portions of the boreal forest that lie well north of established monitoring programmes (Saracco, DeSante & Kaschube 2008; Sauer & Link 2011); however, they are well sampled by migrant banding stations (Leppold & Mulvihill 2011), so an approach that could estimate survival, fecundity and population trajectory as birds pass southward during fall migration would be very useful for population monitoring, and could be applicable to numerous other Holarctic species with extreme northern breeding distributions (Spina 1999; Hussell & Ralph 2005). Model-based fecundity estimates seemed reasonable for both pintails and juncos, but precision was poor given small numbers of same-season recaptures, so I also conducted a simulation study to identify necessary sample sizes for obtaining more precise estimates. This approach provides new opportunities to estimate annual fecundity at local to continental scales and could greatly leverage the utility of existing banding data by allowing investigators to estimate a complete ensemble of vital rates from tagging studies.

## 2 | MODEL AND METHODS

The model developed here assumes that animals are captured after the breeding season has ended using methods that are similarly effective at capturing adults and young of the year, and that captured individuals can be reliably aged at time of marking (e.g., Pyle 1997).

### 2.3 | Model development

A naïve estimator of annual fecundity that ignores differential vulnerability to capture is number of newly marked juvenile females divided by number of newly marked adult females *M*_*jF*_/*M*_*aF*_ (or for sexually monomorphic species, *M*_*j*_/*M*_*a*_). Equation 2 for harvest age ratios can be modified for tag-recovery data as:

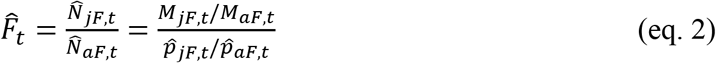

where 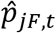 is estimated capture probability for juvenile females in year t, 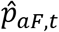 is the equivalent parameter for adult females, and 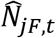 and 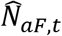 are Horvitz & Thompson (1952) estimators of population size at time of capture. Any appropriate closed-population mark-recapture model could be used to estimate capture probabilities, but given the sparseness of recapture data in my examples, I used Chao’s (1989) estimator, which conditions on the number of individuals captured 1 versus 2 times. Under this model, vulnerability adjusted fecundity 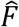 can be estimated as:

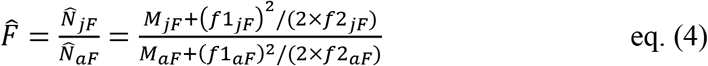

where 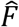 is the estimated age ratio, 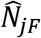 and 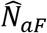 are estimated populations of juvenile and adult females that were available for capture, *M*_*jF*_ and *M*_*aF*_ were the total numbers of juveniles and adults captured and marked (i.e., *M*_*jF*_/*M*_*aF*_ is a naive estimate of fecundity), and *f*1_*jF*_, *f*2_*jF*_, *f*1_*aF*_, and *f*2_*aF*_ were the numbers of juvenile and adult females captured 1 or 2 times, respectively 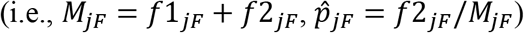. Relative vulnerability 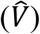 to capture can then be estimated as:

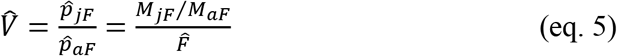

Estimation of age ratios from live recapture data requires a similar set of assumptions as simple closed-population mark-recapture models (i.e. Model M_0_; Otis *et al.* 1978), namely that:1) the population is closed during sampling, 2) animals do not lose marks, 3) all marks are reported on discovery, 4) within age groups, all individuals have the same probability of capture and 5) marking animals does not affect their subsequent catchability. These assumptions have been treated in detail elsewhere (Otis *et al.* 1978), so I focus here on potential violations specific to their application for estimating age ratios. To satisfy the closure assumption, analysts need to select appropriate marking periods for assessing post birth-pulse age structure; choosing intervals after young have become mobile, but before post-breeding dispersal or differential migration have altered age ratios. If data are collected during migration, then ringing operations should include the entire migration period so that capture data are not affected by differential migration of adults versus juveniles (Kelly & Finch 2000). Marker loss is negligible for same-season recaptures, but ironically many North American banders do not report same-season live encounters because the Bird Banding Laboratory historically discouraged such reports (Buckley *et al.* 1998). Homogeneity of capture probabilities among individuals and absence of behavioral response to capture are assumptions that can be accommodated under more elaborate models (Otis *et al.* 1978), but these assumptions are difficult to test with sparse data (Chao 1989).

### 2.2 | Application

Examples used in this paper include northern pintails captured primarily using bait traps on their North American breeding grounds during July through September (Bartzen & Dufour 2017) and dark-eyed juncos captured primarily using mist nets throughout North America during August – October migration (Hussell & Ralph 2005). I used data from 64,201 juvenile and 62,341 adult female northern pintails banded throughout the United States and Canada during 1970-1993 and shot or found dead during the hunting season (1 Sep- 31 Jan of year t+1) 1970-1993 to assess performance of the fecundity model. In addition to the 3841 and 2377 dead recoveries obtained from juveniles and adults during subsequent hunting seasons, there were 90 and 44 live recaptures recorded during the initial banding season. For pintails, annual survey data were available from the Waterfowl Breeding Pair and Habitat Survey (U.S. Fish and Wildlife Service 2017), which indicated a severe population decline during this time period. For dark-eyed juncos, data included 248,939 and 107,998 juveniles and adults banded between 1955 and 2013, but only 121 and 68 dead recoveries and 45 and 15 live encounters during the initial banding season.

I summarized data on number of bandings, dead recoveries and live encounters during the initial trapping period into m-arrays following standard procedures for band-recovery analysis (Brownie *et al.* 1978), except I included an initial column for recaptures (live encounters) during the initial capture period. For juncos, I summarized data in collapsed m-array format recognizing only years since banding (Kéry & Schaub 2012:256) because data were too sparse to consider annual variation in survival or encounter probabilities. Summarized m-arrays and additional details about the data, analysis and JAGS code are provided as supplemental materials.

As an initial template for analysis, I used Seber’s (1971) model for estimating survival (*S*) and reporting rates (*r*) from dead-recovery data, as coded by Kéry & Schaub (2012) for analysis in WinBUGS and further modified for analysis in JAGS 3.3.0 (Plummer 2012) using the jagsUI package in R (Kellner 2015). I first considered models that treated all parameters as constant through time (*S*_*jF*_, *S*_*aF*_, *r*_*jF*_, *r*_*aF*_, *p*_*jF*_, *p*_*aF*_). For pintails, which had more extensive data, I also considered models that included temporal and age-specific variation in survival and recovery (*S*_*j,t*_, *S*_*a,t*_, *r*_*j,t*_, *r*_*a,t*_), but recapture data were too sparse so I treated capture probabilities (*P*_*j*_, *p*_*a*_) as age-specific constants in all models. Time constant parameters were given vague uniform priors on the real scale (i.e. Uniform[0,1]) whereas temporally variable parameters were given vague priors on the logit scale (mean ~ Uniform[−2,2], SD ~ Uniform[0,2]). For pintails, I used an initial 1000 iteration adaptation phase, followed by three Markov chain Monte Carlo (MCMC) chains of 25,000 iterations each, with the first 5000 iterations discarded as burn-in, and retaining every 10^th^ iteration for sampling from the posterior distribution. For juncos, I increased all iterations by 10-fold to accommodate sparse data. Convergence was achieved for all parameters (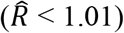 with run times of < 1 minute. Vulnerability (*V*), annual fecundity (*F*_*t*_) and finite population growth (λ_*t*_) were estimated as derived parameters:

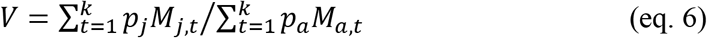

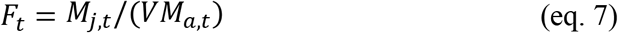

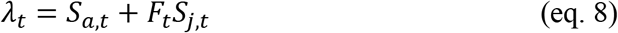

### 2.3 | Simulation specifications

I conducted 1000 24-year simulations representing a data-rich scenario patterned roughly on the northern pintail data; for each simulation I kept S_a_, S_j_ and F constant at 0.60, 0.50 and 0.80, respectively (hence, λ = *S*_*a*_ + *S*_*j*_*F* = 1), but varied number of recoveries and recaptures by using random uniform distributions on r (r_j_ ~ U[0.0001, 0.4]; r_a_ ~ U[0.0001, 0.2]), p (p_a_ ~ U[0.0001, 0.02]) and V (U[0.5, 1.5], with p_j_ = V×p_a_) to produce varying numbers of bandings and live and dead encounters for fixed population sizes of N_j_ = 240 000 and N_a_ = 300 000. In addition to estimates of the mean and standard deviation (SD), I calculated bias, coefficient of variation (CV) and root mean-squared error (RMSE = √[bias^2^ + SD^2^]) for all population and encounter parameters. I compared the accuracy (RMSE) and precision (CV) of these estimates to numbers of juveniles and adults that were banded, recaptured during the first season following banding, or recovered dead during their first or subsequent years to characterize how parameter estimates were affected by variation in quantity of data.

## 3 | RESULTS

### 3.1 | Case studies

Juvenile pintails were more likely to be recaptured than adults (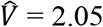, 90% credible interval [CRI]: 1.49 – 2.73), but uncertainty in this parameter translated into large uncertainty in estimates of adjusted fecundity and population growth rate. In the simplest model with no temporal variation, survival and recovery rates, unadjusted age ratios and λ were precisely estimated (CV < 0.1); but recapture rates, vulnerability and adjusted age ratios all had CVs between 0.1 and 0.2 (Table 1). In a fully temporal model, adult survival averaged 0.601 with essentially no annual variation (SD_t_ = 0.002), juvenile survival averaged 0.654 with modest annual variation (SD_t_ = 0.064) and fecundity averaged 0.520 with extensive annual variation (SD_t_ = 0.227). Annual variation in λ_*t*_ was strongly correlated with estimated fecundity (Pearson’s *r* = 0.97), but not with adult or juvenile survival (Fig. 1). Mean annual population growth under both models (time constant: λ = 0.929, 90% CRI: 0.841 – 1.035; time varying: 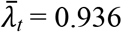, SD_t_ = 0.143) included the estimate derived from survey data (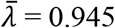, SE = 0.001).

**Table 1:** Estimates of juvenile (juv) and adult (ad) annual survival (*S*), dead recovery (*r*) and live recapture (*p*) probabilities and associated estimates of capture vulnerability (*V*), age ratios at capture (M_j_/M_a_), fecundity (*F*) and finite population growth (λ) for northern pintails and darkeyed juncos under time-constant models.

|  | Northern pintails |  |  | Dark-eyed juncos |  |  |
| --- | --- | --- | --- | --- | --- | --- |
|  | mean | SD | CV | mean | SD | CV |
| S.juv | 0.629 | 0.021 | 0.033 | 0.276 | 0.055 | 0.200 |
| S.ad | 0.613 | 0.0054 | 0.009 | 0.493 | 0.034 | 0.070 |
| r.juv | 0.097 | 0.006 | 0.061 | 0.00045 | 0.00006 | 0.138 |
| r.ad | 0.0412 | 0.0008 | 0.020 | 0.00063 | 0.00008 | 0.122 |
| p.juv | 0.00144 | 0.00015 | 0.105 | 0.00019 | 0.00003 | 0.147 |
| p.ad | 0.00072 | 0.00011 | 0.153 | 0.00015 | 0.00004 | 0.250 |
| V | 2.04 | 0.38 | 0.187 | 1.330 | 0.410 | 0.308 |
| $M_j/M_a$ | 0.992 | 0.0057 | 0.006 | 2.305 | 0.008 | 0.004 |
| F | 0.503 | 0.094 | 0.186 | 1.889 | 0.556 | 0.294 |
| $\lambda$ | 0.929 | 0.060 | 0.064 | 1.014 | 0.191 | 0.189 |

**Fig. 1:**
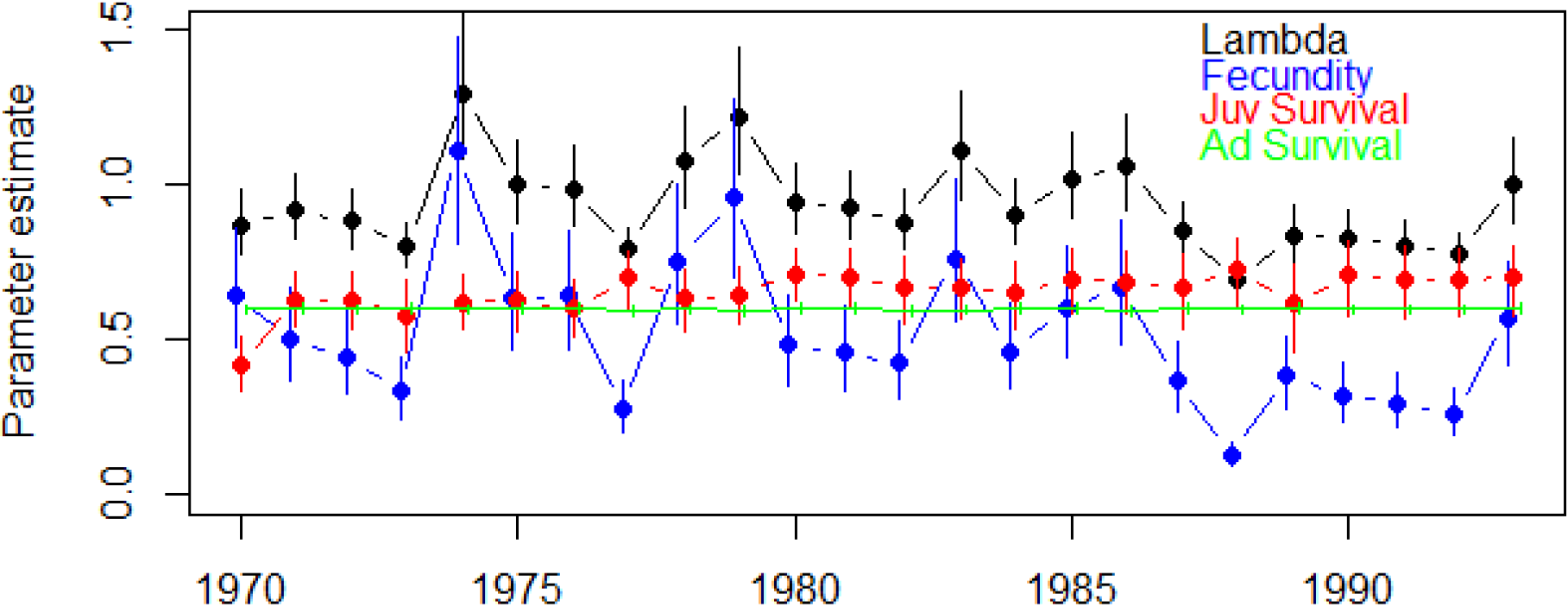
Annual estimates of juvenile survival (JuvS), adult survival (AdS), fecundity (F) and annual population growth (Lambda) for northern pintails during 1970-1993. Fecundity was estimated from age ratios at capture and explained most of the annual variation in lambda.

For juncos, juvenile vulnerability to capture was imprecisely estimated with a credible interval that overlapped 1 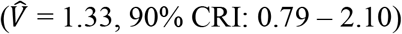. Only unadjusted (raw) age ratios and adult survival were precisely estimated (CV < 0.1), with remaining parameters having CVs exceeding 0.12 (Table 1). Estimated λ was 1.015 (90% CRI: 0.755-1.371), which included the continental estimate based on Breeding Bird Survey data (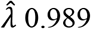, 95% CRI: 0.983-0.995; Sauer & Link 2011).

### 3.2 | Simulations

Precision and accuracy (i.e., lower CV and RMSE, respectively) of juvenile and adult survival and recovery probabilities increased with increasing juvenile, adult, and total recoveries, but these relationships were strongest for adult recoveries (Fig. 1S). Accuracy of vulnerability and adjusted fecundity was most strongly affected by total number of live encounters (Fig. 2). Reasonable estimates (CV < 0.20) of these two parameters required > 300 total recaptures, whereas precise estimates (CV < 0.10) required > 1000 total recaptures. Because estimates of population growth (λ) depended on both survival and fecundity, accuracy of λ estimates were affected by both recoveries and recaptures, but recaptures had the strongest influence.

**Fig 2:**
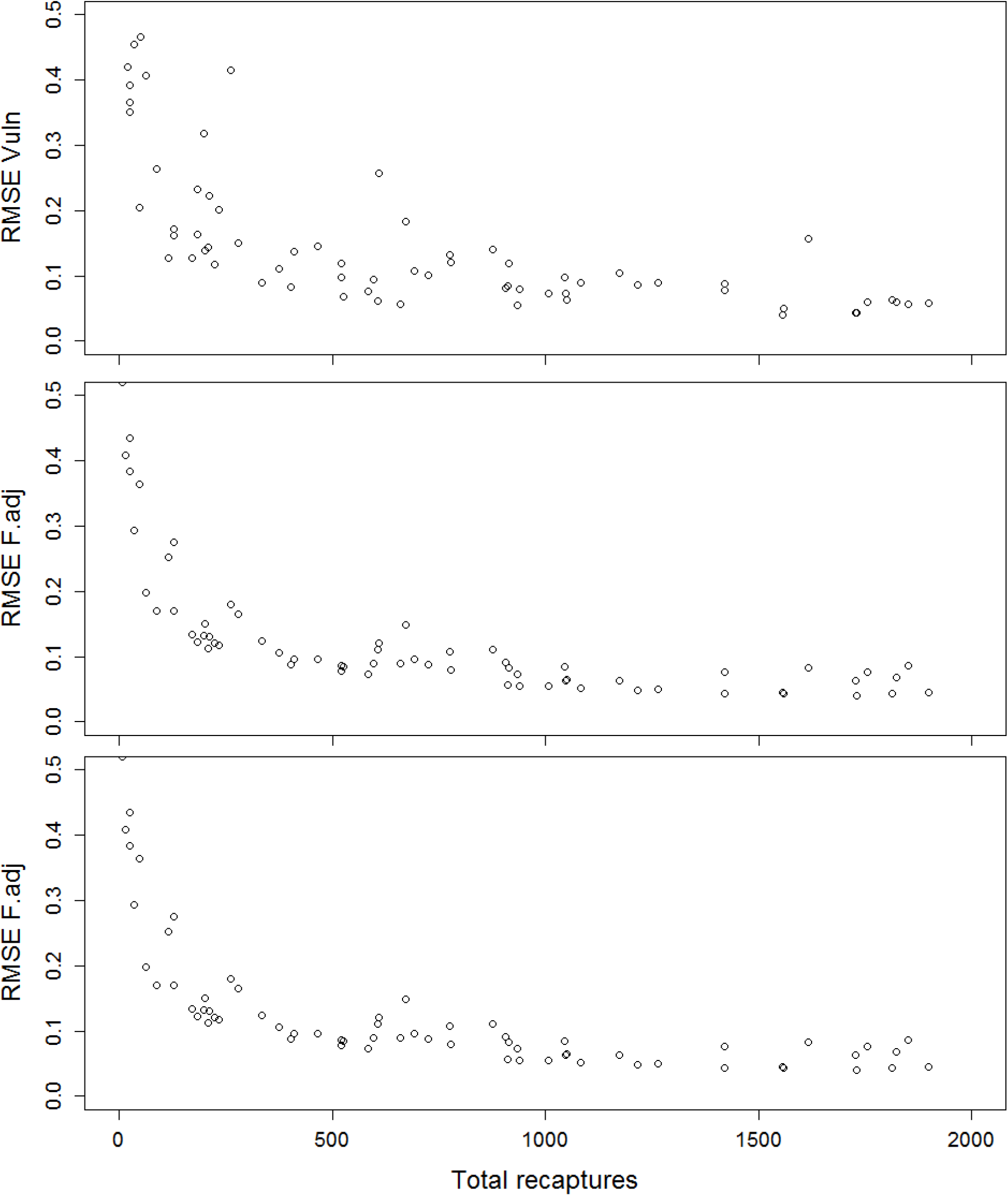
Effect of number of live recaptures (combined juvenile and adult) during the initial marking period on root mean-squared error (RMSE) of vulnerability to capture (Vuln), annual fecundity (F.adj) and population growth rate (lambda).

## 4 | DISCUSSION

Using empirical data on age ratios at capture for northern pintails and dark-eyed juncos, I was able to obtain estimates of annual fecundity by adjusting for vulnerability to capture using live encounters obtained during the original banding season. These fecundity estimates complemented estimates of juvenile and adult survival that analysts have typically obtained from tag-recovery data (Brownie *et al.* 1978; Siriwardena, Baillie & Wilson 1998) and allowed me to construct full life-cycle models that included all of the demographic components of population growth (i.e., λ). For pintails, data were sufficient to estimate annual variation in all three vital rates and these estimates suggested that observed population declines during 1970–1993 were driven primarily by reductions in annual fecundity, which is consistent with other recent studies of historical pintail data (Bartzen & Dufour 2017; Specht & Arnold 2018). For juncos,

Avian ecologists often have access to large-scale count data to assess annual fluctuations in population size (Newson *et al.* 2008; Sauer & Link 2011) and continental banding or ringing programmes can provide similar data on age-specific survival (Francis 1995; Siriwardena, Baillie & Wilson 1998; Saracco *et al.* 2010), but fecundity data are often lacking (Ahrestani *et al.* 2017). To assess fecundity, population modelers have used age ratios at harvest from hunted species (Péron & Koons 2012), fledgling counts from citizen-scientist nest-record programmes (Robinson, Morrison & Baillie 2014), data from small-scale nesting studies (Weegman *et al.* 2017) and reverse-time mark-recapture models (Saracco, DeSante & Kaschube 2008), but age ratios at capture could provide an alternative or complementary data stream to assess spatiotemporal variation in fecundity (Mazerolle *et al.* 2005; Ross *et al.* 2017; Specht & Arnold 2018). In the absence of live recapture data, vulnerability to capture (*V*) could be estimated in an integrated population modeling (IPM) framework (Ahrestani *et al.* 2017), assuming that auxiliary population count data were available and that there were no confounding influences of immigration or emigration:

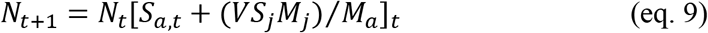

If marking efforts occur at the end of the breeding season, but before post-breeding dispersal or migration, then age ratios at marking measure local reproductive success and spatially extensive marking data have the potential to measure regional variation in fecundity and identify ecological or anthropogenic drivers of this variation (Specht & Arnold 2018). However, researchers must have a thorough understanding of breeding and movement phenology to select appropriate intervals and spatial scales for data analysis, thereby assuring that age ratios are not affected by ongoing breeding efforts or early dispersal or migration by one age class versus another (Andres, Browne & Brann 2005). Variation in age ratios at capture could also be due to age-related variation in local habitat use on the breeding grounds, especially if capture efforts are not randomly distributed among potential habitats. Treating individual capture sites as random effects could potentially control for some of this location-specific variation (Specht & Arnold 2018) and testing for seasonal trends in age ratios could help identify ongoing breeding or differential movements. Age ratios might also be affected by capture methods, if juveniles are more (or less) vulnerable to capture by widely employed capture methods. In North America, relatively few capture methods are uniquely coded at time of banding (https://www.pwrc.usgs.gov/BBL/MANUAL/summary.cfm), but European ringing schemes record a wide diversity of capture methods and lure types (https://euring.org/files/documents/E2000PLUSExchangeCodev1161.pdf), thereby allowing for a thorough investigation of heterogeneity in age ratios induced by capture methodology.

In the northern hemisphere, many birds are banded or ringed during autumn as they migrate from northern hemisphere breeding sites to equatorial or southern hemisphere wintering sites (Spina 1999; Hussell & Ralph 2005). Such marking programmes have the potential to assess continental-level productivity, but meeting the closure assumption seems much more difficult in this situation (Hochachka & Fiedler 2008). Nichols *et al.* (2009) partitioned detection probability from count surveys into four conditional components, and a similar hierarchy could be extended to capture probabilities. First, choice of marking sites could affect age ratios at capture if juveniles and adults have different migration routes (Ralph 1978). Second, differential timing of migration could affect age ratios at capture (Andres, Browne & Brann 2005), especially if one age class exhibits a more prolonged migration and capture efforts are limited to periods of peak migration. Third, age-related capture probability could be affected by differences in stopover durations; for example, if juveniles spend more time “refueling” at migrational stopover sites they would be more vulnerable to capture (Rguibi-Idrissi, Julliard & Bairlein 2003), especially if permanent marking sites are concentrated at migrational stopover sites. Finally, because juveniles are more naïve, they may be more vulnerable to capture by standard trapping methods (Rguibi-Idrissi, Julliard & Bairlein 2003), even if locations and timing were unbiased.

Probably the biggest limitation to employing the fecundity estimation approach developed herein is the paucity of same-season live-encounter data for estimating vulnerability to initial capture. With sufficient recapture data, many of these assumptions could be tested, and some ringing stations have sufficient in-house data to estimate capture vulnerability (e.g., Hochachka & Fiedler 2008). During routine duck banding operations in Alberta, Canada, approximately 54% of 33,552 ducks captured for banding over a 3-year period were same-season recaptures (Dieter, Murano & Galster 2009), but banding crews have not been encouraged to collect and report these data. North American banders were historically dissuaded from reporting same-station live encounters, and hence live encounter data are limiting for historical analyses, although this shortcoming has been recently corrected (Smith 2013) and many North American banders have begun submitting large amounts of recapture data (D. Bystrak, Patuxent Wildlife Research Center, *pers. comm*.). In Europe, many national ringing programmes failed to keep records of numbers of bands deployed and focused primarily on banding known-age juveniles, but this shortcoming was recognized in the mid-1980s (e.g., Anderson, Burnham & White 1985) and ringing schemes have since expanded to include adults, and historical summaries of ring deployment have since been compiled for many European countries going back to 1975 (https://euring.org/data-and-codes/ringing-totals). Bird ringers need to be made aware of the value of live encounters, even those from the same location and banding season, and national banding programmes need to be made aware of the value of collecting and archiving such data. The ability to estimate fecundity from age ratios at the time of marking greatly enhances the utility of continental ringing programmes, because it allows important vital rates to be estimated as markers are deployed, while investigators wait for encounter data to accumulate.

## ACKNOWLEDGEMENTS

I am grateful to the countless individuals who deployed bands on pintails and juncos, and to all the hunters and citizens who reported encounters to the Bird Banding Laboratory. Hannah Specht provided helpful feedback on an earlier draft of the manuscript.

## DATA ACCESSIBLITY

Data and code used for analysis and simulations will be made publicly available.

---

**Fig S1:**
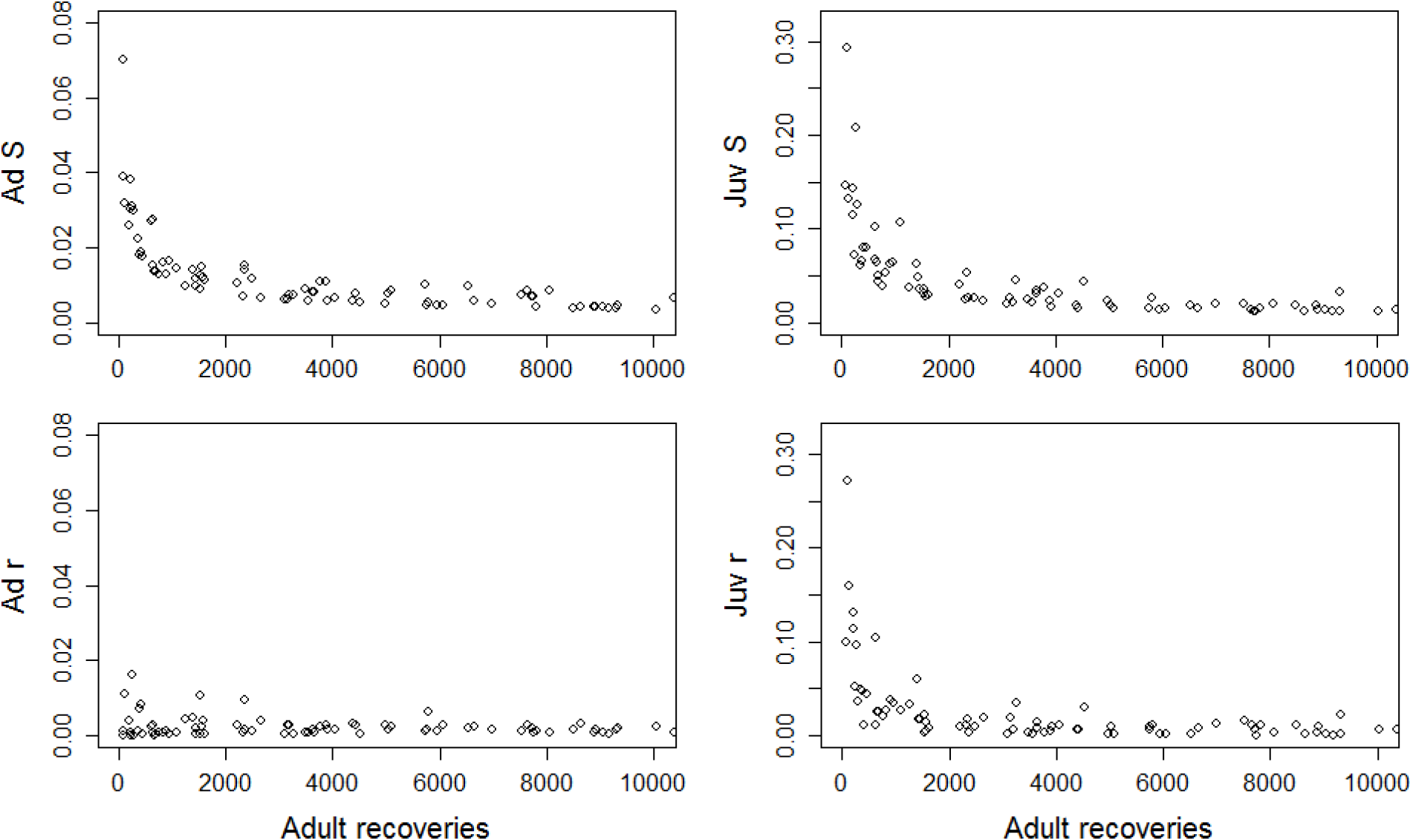
Effect of number of adult recoveries on accuracy (root mean-squared error) of juvenile and adult survival and reporting rates from Seber recovery models. Note the RMSE scale for juveniles (right column) is 4-fold higher than for adults. In addition to variation in adult recoveries, juvenile recovery probabilities also varied randomly from 0 to 0.4 across all simulations, contributing to excess variation.

